# Accelerated prime-and-trap vaccine regimen in mice using repRNA-based CSP malaria vaccine

**DOI:** 10.1101/2023.05.23.541932

**Authors:** Zachary MacMillen, Kiara Hatzakis, Adrian Simpson, Melanie J. Shears, Felicia Watson, Jesse H. Erasmus, Amit P. Khandhar, Brandon Wilder, Sean C. Murphy, Steven G. Reed, James W. Davie, Marion Avril

## Abstract

Malaria, caused by *Plasmodium* parasites, remains one of the most devastating infectious diseases worldwide, despite control efforts that have lowered morbidity and mortality. The only *P. falciparum* vaccine candidates to show field efficacy are those targeting the asymptomatic pre-erythrocytic (PE) stages of infection. The subunit (SU) RTS,S/AS01 vaccine, the only licensed malaria vaccine to date, is only modestly effective against clinical malaria. Both RTS,S/AS01 and the SU R21 vaccine candidate target the PE sporozoite (spz) circumsporozoite (CS) protein. These candidates elicit high-titer antibodies that provide short-term protection from disease, but do not induce the liver-resident memory CD8^+^ T cells (Trm) that confer strong PE immunity and long-term protection. In contrast, whole-organism (WO) vaccines, employing for example radiation-attenuated spz (RAS), elicit both high antibody titers and Trm, and have achieved high levels of sterilizing protection. However, they require multiple intravenous (IV) doses, which must be administered at intervals of several weeks, complicating mass administration in the field. Moreover, the quantities of spz required present production difficulties. To reduce reliance on WO while maintaining protection via both antibodies and Trm responses, we have developed an accelerated vaccination regimen that combines two distinct agents in a prime-and-trap strategy. While the priming dose is a self-replicating RNA encoding *P. yoelii* CS protein, delivered via an advanced cationic nanocarrier (LION^TM^), the trapping dose consists of WO RAS. This accelerated regime confers sterile protection in the *P. yoelii* mouse model of malaria. Our approach presents a clear path to late-stage preclinical and clinical testing of dose-sparing, same-day regimens that can confer sterilizing protection against malaria.

## INTRODUCTION

Despite control efforts that have reduced morbidity and mortality, malaria remains a major burden globally, with deaths and cases rising in recent years^1^. Research efforts have led to the development of several vaccine approaches, ranging from whole organism (WO) parasite-based vaccines to subunit (SU) vaccines delivered in protein-in-adjuvant platforms or in viral or bacterial vectors^2^. The SU vaccine RTS,S/AS01E, the only malaria vaccine approved by the World Health Organization (WHO), is composed of virus-like particles consisting of hepatitis B virus surface antigen (HBsAg) combined with a fragment of *Plasmodium falciparum* (Pf) circumsporozoite (CS) protein. While approval of the RTS,S vaccine represents a significant step forward, it is unlikely to lead to eradication of malaria, given its modest efficacy, which declines further over years ^3–7^. More recently, the SU R21-MatrixM vaccine, which uses the same CS protein as RTS,S but with an increased ratio of CS to HBsAg, has also exhibited significant efficacy in a seasonal malaria setting. Early results of a clinical trial of R21-MatrixM among children living in an endemic area showed 77% efficacy over a 12-month period in a seasonal setting ^8^. Unlike RTS,S, the R21 vaccine is practical to produce at scale. However, the utility of both vaccines is severely limited by the requirement for 3-4 doses plus seasonal boosters to achieve the strategic goal of the WHO of 75% protective efficacy against clinical malaria.

Development of SU vaccines against *Plasmodium* is challenging, due to the parasite’s large genome and complex multi-host life cycle. Eliminating the parasite at the clinically-silent PE stage is a particularly attractive goal, as it would prevent erythrocytic infection and thus halt both disease and transmission^7,8^. One of the primary candidates for PE vaccine development is the CS protein, which is expressed by infectious spz and is required for parasite motility and hepatocyte invasion. CS is composed of an N-terminal region that binds heparin sulfate proteoglycans (RI), an immunodominant central repeat region of four amino acids (NANP) that is the target of neutralizing antibodies, and a GPI-anchored C-terminal region containing a thrombospondin-like domain (RII) and T-cell epitopes ^5,9,10^. Numerous vaccine platforms using CS-based virus-like particles, self-assembling protein nanoparticles, viral or bacterial vectors encoding CS, or even high-affinity monoclonal anti-CS antibodies used as therapeutics, have been evaluated as malaria interventions, but have not met the critical benchmarks ^10–17^. Other strategies focus on delivery of WO vaccines, which generate a diverse immune response that recruits both antibodies and T cells targeting numerous antigens. For example, RAS immunization can elicit high antibody titers, central memory and effector memory CD8+ T cells (Tcm and Tem respectively), and resident (non-circulating) memory T cells (Trm)^18^. These Tem and Trm can mediate protection in WO-immunized mice by eliminating infected hepatocytes and conferring sterilizing immunity ^19–21^. Thus, one hypothesis is that robust PE immunity may depend on generation of liver-resident Trm. Protection has been achieved in both animal models and humans using repeated immunizations with WO RAS. Vaccination strategies employing WO spz (RAS, genetically-attenuated parasites (GAP), or chemoprophylaxis combined with administration of Pf spz (known as CPS)) have successfully protected malaria-naïve volunteers^22–26^. The protection appears to be mediated by a combination of liver-resident Trm and antibodies^27^. However, the WO RAS vaccination strategy was less effective in malaria-endemic regions, confirming the need for more robust approaches ^28–31^.

The vaccination method known as “prime-and-trap” is intended to generate cellular immunity, by inducing liver Trm-mediated protection, as well as humoral immunity, with the help of CD4+ T cells and antibody production by B cells. Previous vaccination studies have assessed both homologous and heterologous prime-boost and prime-target approaches. These include, for example, viral-vectored vaccines harboring *Plasmodium* antigens such as Fowlpox Virus 9 (FP9), chimpanzee adenovirus 63 (ChAd63) combined with the attenuated vaccinia virus (Ankara, MVA)^11,32^, and a two-dose “prime-and-target” method (so named to avoid confusion with the TRAP antigen employed in the formulation) using a human adenovirus type 5 (AdHu5) prime followed by a boost using adeno-associated virus stereotype 1 (AAV1) ^33^. These approaches have been shown to induce both specific, functional antibodies and high numbers of Trm at the time of infection, but none of these has achieved the strong safety profile of WO vaccination. Other “prime-and-trap” approaches combine a priming dose of a nucleic acid with a heterologous trapping dose of WO spz that naturally home to the liver ^34,35^. The resulting liver Trm are positioned to respond effectively to liver-stage parasites, leading to sterile protection. Prime-and-trap vaccination using a nucleic-acid prime and WO RAS trap in BALB/cJ mice completely protected against WT rodent malaria (*Py*) spz challenge ^34^. Importantly, this approach required only one dose of WO spz ^35^. However, the prime-and-trap vaccines developed to date have generally relied on DNA vaccination delivered by gene gun^34^, and there is no gene gun delivery device currently approved for clinical use.

Lipid nanoparticles (LNPs) are a novel and highly effective means of delivering ribonucleic acid (RNA) vaccines. Antigen-encoding messenger RNA (mRNA) emerged as an optimal strategy for vaccination during the COVID-19 pandemic: these vaccines express antigen upon delivery into tissue, stimulate the innate immune system, induce cellular immunity, and eliminate the need to support large-scale protein antigen production. When PfCSP mRNA was combined with a lipid nanoparticle (LNP) and tested in mice, a three-dose regimen achieved up to 60% protection in a Pb(ANKA)-PfCSP challenge model in Balb/c and C57Bl6 ^36^. Long-term protection was not observed, again highlighting the limitations of SU-only approaches. More recently, mRNAs encoding the PfCSP and Pfs25 proteins were formulated with LNP and used to elicit functionally effective immune responses in mice to both antigens. This formulation protected against spz challenge and reduced Pf transmission to mosquitoes after multiple immunizations ^37^. However, clinical trials with mRNA vaccines formulated with traditional LNPs ^38–40^ have encountered challenges such as LNP/mRNA reactogenicity upon injection, biodistribution to multiple organs^41^,^42^, and instability during prolonged storage ^43^.

Self-replicating or replicon RNA (repRNA) is a new and promising alternative to mRNA vaccines. Once in cells, repRNA initiates biosynthesis of antigen-encoding mRNA, raising and prolonging antigen expression and enhancing humoral and cellular immune responses^44^. Based on recent studies ^44–46^, repRNA systems formulated in an oil-in-water emulsion nanocarrier exhibit greater efficacy at lower doses than mRNA vaccine platforms. Indeed, repRNA encoding Zika virus antigens can completely protect mice against a lethal ZIKV challenge with a single dose (10 ng RNA) ^45^. Moreover, repRNA elicits more robust immune responses after a single dose than conventional mRNA formulations, offering an attractive approach for emerging infectious diseases such as dengue ^46^, Zika^45^, and SARS-CoV-2/COVID-19 ^44^.

While repRNA is outstanding for promoting antigen expression, the *in vivo* instability of RNA and the requirement for transport through lipid bilayers have stimulated development of novel vehicles for intracellular RNA delivery. As an alternative to LNP-encapsulated mRNA, repRNA was formulated with LION™, a cationic oil-in-water nanoparticle emulsion designed to enhance vaccine stability, delivery, and immunogenicity ^44^. In response to the SARS-CoV-2/COVID-19 pandemic, Erasmus *et. al*. developed a repRNA vaccine candidate, repRNA-CoV2S, encoding the SARS-CoV-2 spike (S) protein formulated with LION. This LION/repRNA-CoV2S construct elicits robust anti-SARS-CoV-2 spike-protein IgG antibody isotypes in mice, indicative of a Type-1 T-helper response^44^. Importantly, a single-dose administration in non-human primates (NHP) elicited antibody responses that potently neutralized SARS-CoV-2^44^. The LION formulation produced under cGMP has undergone toxicology evaluation in rabbits in conjunction with repRNA for SARS-CoV-2 and is currently in Phase I/II human clinical trials in South Korea ^47^, Brazil (clinicalTrials.gov Identifier: NCT05542693), and the US (clinicalTrials.gov Identifier: NCT05132907). This LION-repRNA-CoV2S vaccine, as a lyophilized drug product, has recently received Emergency Use Authorization by regulators in India as the first self-amplifying RNA vaccine ever approved for human use ^48^, with potential advantages of lower cost and higher stability that should facilitate global distribution.

Here, we leverage this LION/repRNA technology to generate a two-dose, same-day prime-and-trap vaccine against malaria in mice. This was achieved by intramuscular (IM) priming with repRNA encoding full-length CS of *Plasmodium yoelii* (Py) (repRNA-PyCS) formulated in the LION nanoparticle carrier, followed by an IV injection of WO RAS as the trapping dose. This two-component, same-day regimen conferred sterile protection in mice and engaged both humoral and cellular arms of the immune system, bringing us closer to a single-visit, highly effective malaria vaccine.

## RESULTS

### repRNA-CS vaccine formulation and prime-boost immunogenicity in BALB/cJ mice

Using the attenuated Venezuelan equine encephalitis (VEE) virus TC-83 strain^49^, we incorporated the coding sequence of the full-length CS protein from Py into the alphavirus expression vector to create a repRNA malaria vaccine. The coding sequences of the full-length CS protein from Pf and *Plasmodium vivax* (Pv) were incorporated into the same expression vector as controls (Supplemental Figure 1A). After RNA transcription and capping, expression of repRNA-PyCS, -PvCS, and -PfCS was verified by denaturing gel electrophoresis (Supplemental Figure 1B). The vectors were then transfected into mammalian cells for validation. Western-blot analysis showed expression of CS proteins at higher apparent molecular weights (MW) than expected (observed vs expected: PyCS ∼99Kd vs 44.7Kd, PfCS ∼70Kd vs 43.4Kd, PvCS ∼60Kd vs 36.9Kd, respectively), probably due to glycosylation that impeded protein migration into the gel (Figure 1A, Supplemental Figure 1C).

**Figure 1.**
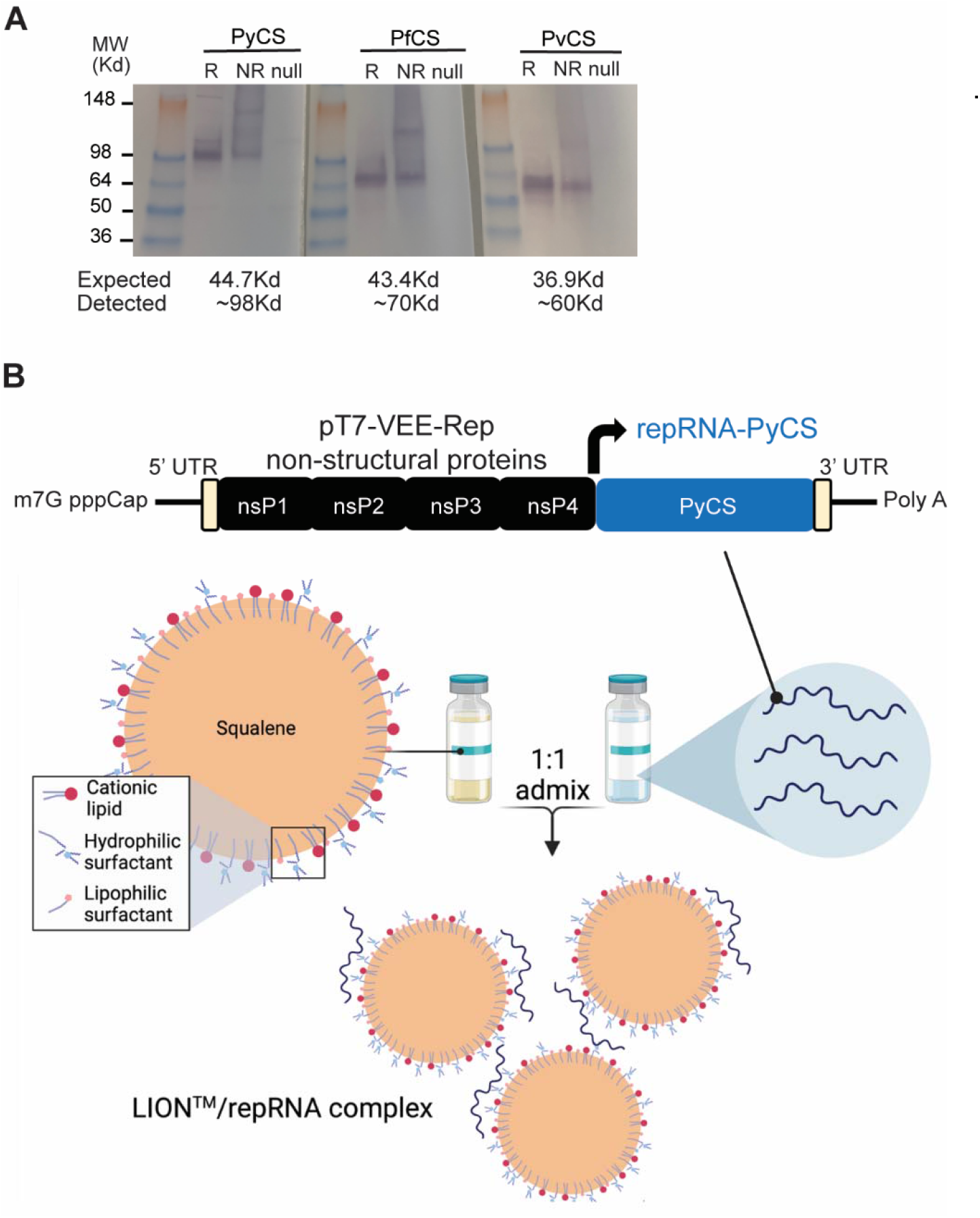
LION/repRNA-CS vaccine design. A) After RNA transcription and capping, repRNA-PyCS or -PfCS or -PvCS was transfected into BHK cells and 24 to 48 hours later, the lysate (R, reduced, NR, non-reduced) of transfected cells or null transfection used as control were analyzed by Western blot, using rabbit polyclonal antibodies for immunodetection. B) Graphic representation of replicon repRNA-PyCS and LION formulation that was used for immunization after mixing

The repRNA-CS vaccines were formulated with the novel LION emulsion (Figure 1B) ^44^. Unlike current mRNA vaccines, the LION/repRNA vaccine platform utilizes an admixture formulation of LION, a highly stable cationic (DOTAP) squalene emulsion embedded in a hydrophobic oil phase (Figure 1B) that can be manufactured independently of the RNA component and combined with the repRNA construct in a simple 1:1 (v/v) mixing step 30 minutes prior to immunization. To determine the immunogenicity of homologous prime-boost LION/repRNA-CS vaccination with single or dual CS antigens, mice were immunized with an IM prime of 5 µg of LION/repRNA, followed by a homologous boost 14 days later (Supplemental Figure 2A). The mice received a LION/repRNA vaccine with either one of the antigens (PyCS or PfCS; 5 μg per antigen) or a combination of two antigens (PyCS and PfCS, or PfCS and repRNA-GFP control; 2.5 μg per antigen). A repRNA-GFP construct was used as a non-malaria-coding antigen control. Final bleeds were collected three weeks post-boost and immune responses were analyzed by ELISA against the corresponding CS tandem-repeat region. Humoral immune responses against corresponding CS antigens as evaluated by ELISA exhibited little to no cross-reactivity against heterologous CS (Supplemental Figure 2B). All mice seroconverted after immunization. The strength of CS-specific responses to the two-antigen-based vaccines was slightly lower than to the single-CS-antigen vaccines (p=0.0286 for PyCS compared to PyCS+PfCS; p=0.7302 for PfCS compared to PyCS+PfCS; Supplemental Figure 2B), possibly because of the reduced dose per antigen in the former. Nevertheless, both 5μg and 2.5μg doses of the prime-boost vaccines were highly immunogenic and demonstrated the feasibility of using lower dosages of two-antigen-based vaccines (Supplemental Figure 2C). These results show that the LION/repRNA-CS platform can induce antibody responses against malaria antigens alone or in combination and suggest that any humoral immune interference occurring when two antigens are mixed in the vaccine is minimal.

### Immunogenicity and efficacy of a two-dose prime-boost repRNA-PyCSP vaccine in BALB/cJ mice

To confirm antibody responses to homologous prime-boost LION/repRNA-PyCS vaccination in BALB/cJ, 15 mice were immunized with IM injections 14 days apart (Figure 2A, Supplemental Table 1). The LION/repRNA-PfCS and LION/repRNA-PvCS vaccines were used as separate controls in cohorts of 7 mice each. Antibody responses were evaluated by ELISA following the prime (day 13) and boost (day 29) against peptides containing their corresponding CS tandem-repeat sequences (Figure 2B). The anti-PyCS total IgG antibody levels in primed mice (D13, p=0.001, Figure 2B) were substantially higher than in naïve mice and were enhanced by a second dose of LION/repRNA-PyCS (day 29, p=0.028, Figure 2B) with slight cross-reactivity observed against the PfCS epitope (Figure 2B).

**Figure 2.**
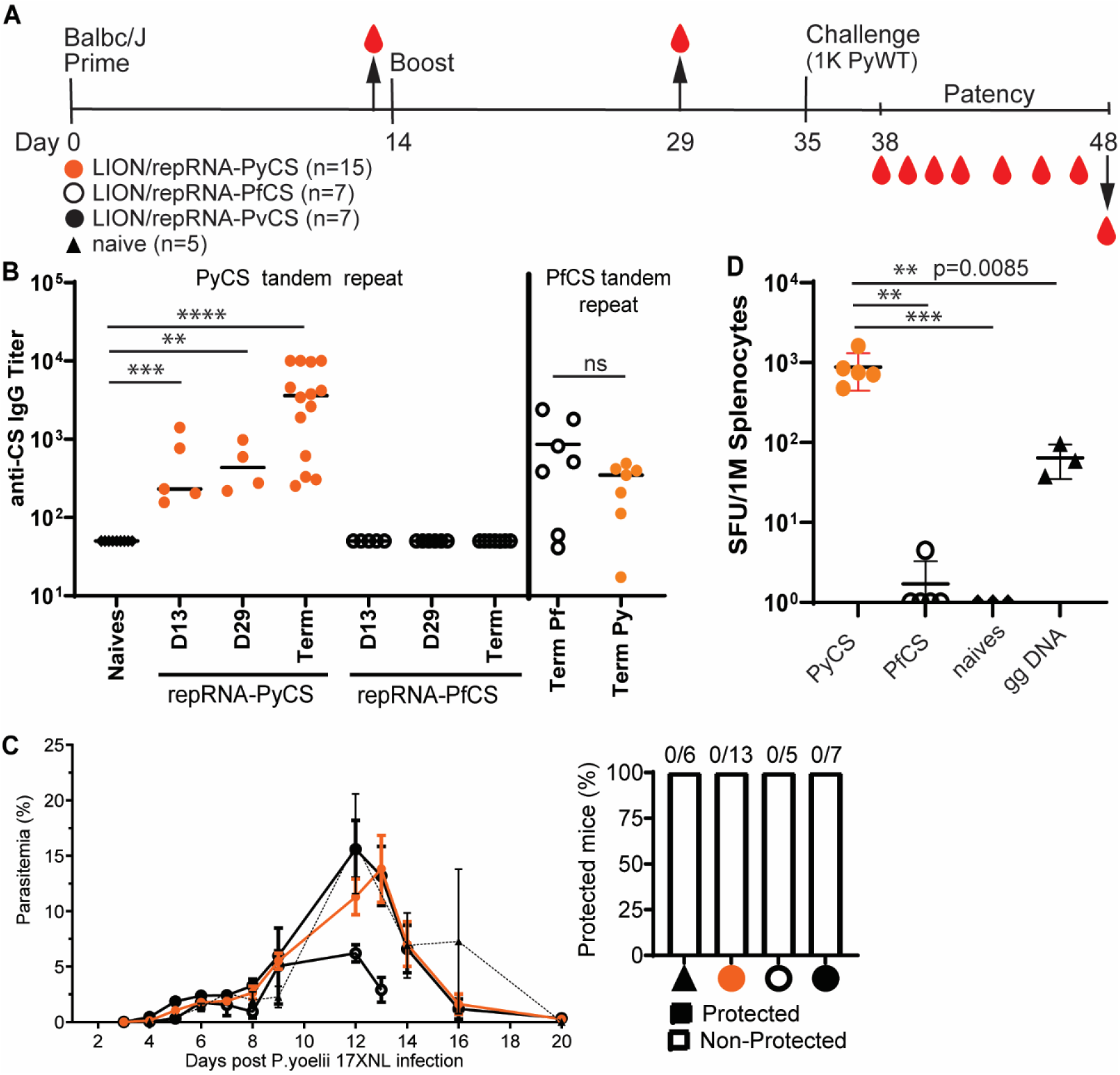
Immunogenicity and efficacy of a prime-boost vaccine with repRNA encoding either PyCS (orange circle), PfCS (opened circle) or PvCS (black circle) formulated with LION. A) BALB/cJ mice immunization schedule. Mice were injected with 5ug of LION/repRNA-PyCS, -PfCS, or -PvCS in a 2-week interval prime-boost regimen. B) Mouse sera were collected at post-prime (day 13), post-boost (day 29), and final bleeds post-challenge (day 48, i.e., terminal bleed). PyCS and PfCS antibody responses were determined by PyCS or PfCS repeat region peptide titration ELISA, respectively. C) Parasitemia and protection post-challenge of the three prime-boost cohorts. Number of mice per cohort indicated above bar graph. Naïve cohort indicated with a triangle symbol and dash line. D) IFNγ ELISPOT of CS-specific T cells four weeks after a single prime injection of LION/repRNA-PyCS. Cohorts receiving gene gun DNA encoding PyCS (ggDNA), LION/repRNA-PfCS, or no immunization were used as controls. The n value represents total number of mice tested per cohort, in two to three independent assays. Each data point represents an individual mouse and the bar represents the group mean. Asterisks represent significance as determined by the non-parametric two-tailed Mann–Whitney two-tailed test (*p=0.05, **p=0.01, ***p=0.001, ****p<0.0001).

To assess efficacy, subsets of mice from each of the LION/repRNA-PyCS, LION/repRNA-PfCS, and LION/repRNA-PvCS cohorts were challenged 3 weeks post-boost with an IV injection of 1000 wildtype (WT) Py 17XNL spz freshly dissected from mosquito salivary glands (Supplemental Table 1). This prime-boost of repRNA-PyCS vaccination alone did not protect against Py wild-type spz challenge in mice (Figure 2C). However, ELISA at the endpoint (post-challenge, noted as “term” at day 48) showed that the antibody levels were recalled by the challenge dose of WT spz (p<0.0001 relative to naives, Figure 2B). These repRNA-PyCS data show the consistency of the antibody responses induced by this repRNA platform, though full protection was not conferred by this homologous approach.

### Superior T cell immunogenicity of repRNA-PyCS over gene-gun DNA priming in BALB/cJ mice

To assess T-cell responses to LION/repRNA-PyCS and evaluate this candidate as a potential priming dose in prime-and-trap vaccination, additional cohorts of BALB/cJ mice were immunized with a single IM injection and responses were assessed by splenocyte IFNγ ELISPOTs four weeks later (Figure 2D). Responses were compared to splenocytes from mice immunized with a single repRNA-PfCS dose or DNA encoding PyCS administered by gene gun (gg DNA-PyCS), as used for priming in the first-generation prime-and-trap vaccine using RAS ^34,35^. As expected, naïve mice and mice primed with the control LION/repRNA-PfCS did not recognize the PyCS epitope, as assessed by ELISPOT. In comparison, the LION/repRNA-PyCS vaccine elicited over 10-fold more IFNγ-producing T cells than gene-gun DNA-PyCS vaccination (p=0.0085, Figure 2D), indicating robust priming of CD8+ T cells following a single LION/repRNA-PyCS injection.

### Immunogenicity of the accelerated prime-and-trap repRNA-PyCS-RAS vaccination in BALB/cJ mice

To assess LION/repRNA-PyCS as the priming dose in prime-and-trap vaccination, cohorts of BALB/cJ mice were immunized with LION/repRNA-PyCS followed by PyRAS, along with control cohorts receiving two doses of homologous prime-boost repRNA-PyCS, or PyRAS trap-only (consisting either of an irrelevant repRNA-PfCS control followed by PyRAS or a single dose of PyRAS alone). Naïve animals were used as additional controls. Previous prime-and-trap vaccine studies ^34,50^ employed a 28-day interval between doses and utilized 20,000 to 50,000 RAS for the trapping dose. To investigate whether the schedule could be accelerated with the LION/repRNA prime and 25,000 PyRAS as the trapping dose, mice were primed IM with repRNA-PyCS (1 μg or 5 μg) 14 or 5 days prior to a PyRAS trapping dose (Figure 3A).

**Figure 3.**
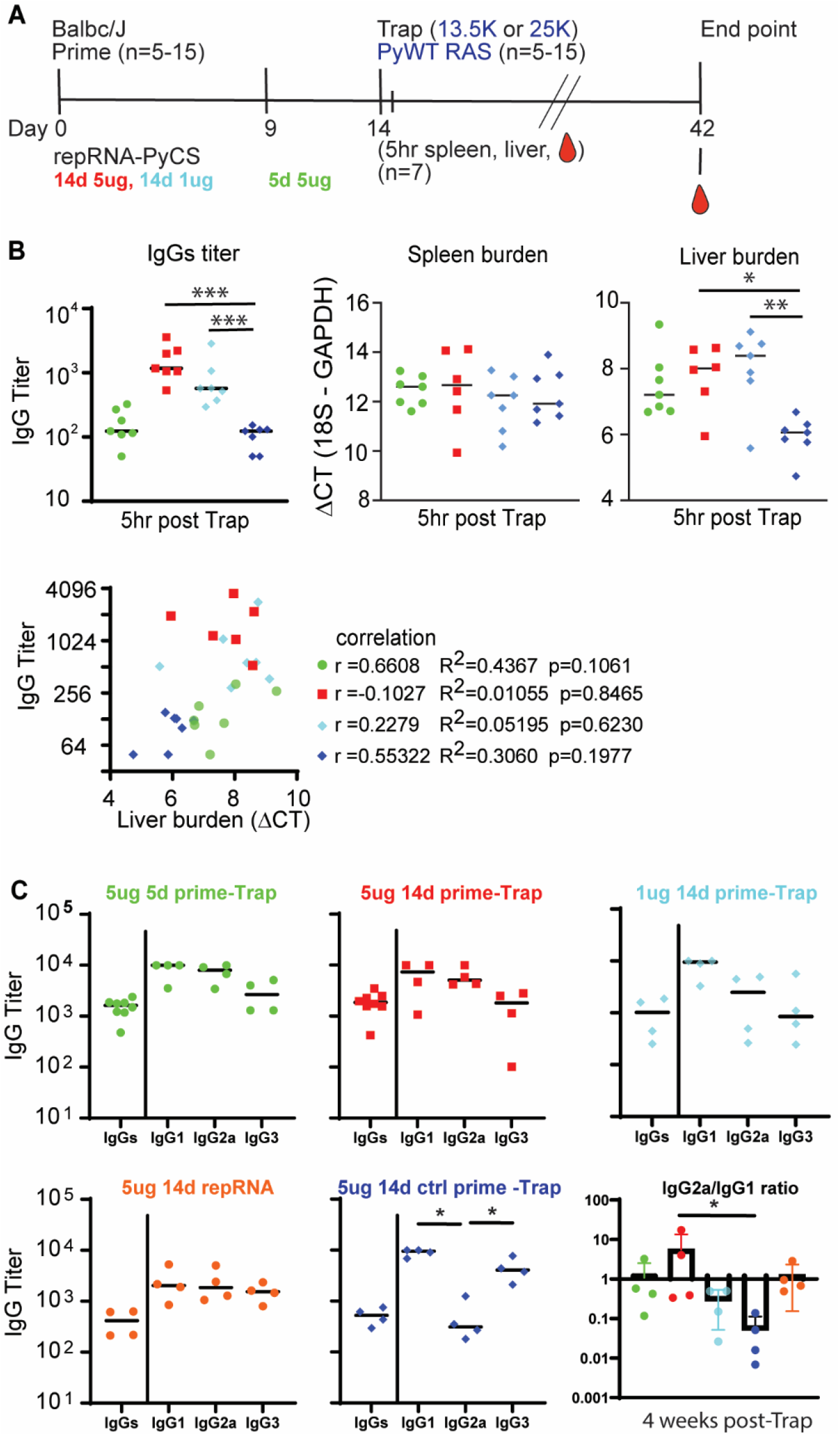
Immunogenicity of accelerated prime-and-trap immunization regimens in BALB/cJ mice. A) 5ug 5-day (green data) or 5ug 14-day (red data) or 1ug 14-day regimen of repRNA-PyCS (light blue data) prime followed by trap dose of 25,000 RAS, were tested in mice. Control cohorts are prime-boost repRNA-PyCS (orange data) or trap cohort (repRNA-PfCS +RAS, dark blue data). A) Immunization schedule. B) Spleen, liver, and serum from seven mice per cohort were harvested 5 hours post-trap injection. Total IgG titer was analyzed by ELISA, while spleen and liver parasite burden were analyzed by qPCR quantification. Spleen and liver burden qPCR was evaluated using one-way ANOVA followed by Kruskal-Wallis test and Dunn’s multiple comparisons test (* p<0.05, ** p<0.005). A lower ΔCT represents a high parasite burden. Pearson correlation comparing the IgG titer and the liver burden for the four vaccine group, For visualization purpose, all cohorts were combined into the same graph, but each correlation coefficient was computed individually for each cohort. C) Final-bleed sera were collected at endpoint, 4 weeks post-trap (day 42) to evaluate total IgG titer and IgG1, IgG2a, IgG3 subclasses for each cohort by ELISA. Ratio of IgG2a/IgG1 is indicated in bar graph. All others statistical analyses were performed using a Mann-Whitney test. The n value represents total number of mice tested per cohort in two independent assays. Each data point represents an individual mouse, and the bar represents the group mean.

To assess CS-specific whole IgG levels and parasite burden post-trapping, we collected sera, spleens, and livers from seven mice of each cohort within 5-6 hours after the trapping dose. As expected, mice immunized with either 14-day regimen (1 μg or 5 μg) had higher antibody titers than mice immunized with a 5-day regimen of repRNA-PyCS (trapped with RAS) and the irrelevant repRNA-PfCS regimen-with-RAS cohort (Figure 3B). No difference was observed among the four cohorts in parasite burden in the spleen, whereas the repRNA-PfCS-plus-RAS cohort and the 5-day cohort exhibited a trending correlation between parasite burden in the liver and high levels of IgG (Figure 3B). These results suggest that specific anti-PyCS IgG generated beyond 14 days post-prime targeted the WO spz from the trapping dose, reducing their distribution to the liver target.

Next, to determine if a balanced or skewed IgG subclass response was induced by repRNA-PyCS, we measured circulating IgG subclasses four weeks post-trapping dose in a separate cohort of mice, using CS peptide ELISA (Figure 3C). There was no significant difference between the IgG1, IgG2a, and IgG3 levels, or in the IgG2a/IgG1 ratio between cohorts immunized with repRNA-PyCS as the priming dose, indicating that the two-dose regimen induces a balanced Th1/Th2 antibody response. However, the cohort receiving RAS alone as the trapping dose (i.e, 14-day repRNA-PfCS and RAS, Figure 3C, dark-blue data) exhibited a skewed Th2-type humoral immune response. Our results implicate IgG2a in addition to the IgG1 and IgG3 subclasses as significant components of the humoral response, as opposed to the narrow IgG1-biased response observed after administration of a WO-based vaccine such as RAS. Additional experiments are ongoing to evaluate the relationships of these protective responses in mice and functional antibodies.

Finally, to assess CD8^+^ T-cell responses to LION/repRNA-PyCS prime-and-trap vaccination, mice were immunized under various regimens (5 μg vs 1 μg as priming dose in a prime-and-trap 14-day vs 5-day regimen; 5 μg prime-boost or 25,000 spz trap-alone regimen) as described above (Figure 4A). Livers were collected 28 days after the last immunization in two independent experiments, and lymphocytes were isolated and stained for flow cytometry as described previously ^34^. Total CD8^+^ T cells and activated CD8^+^ T cells (CD44^hi^ CD62L^lo^) in the livers of prime-and-trap (5 μg 5-day, 5 μg 14-day, 1 μg 14-day) or trap-only (i.e., control prime-and-trap (5 μg 14-day) or 25,000 RAS) immunized mice were significantly higher than in the homologous prime-boost repRNA-PyCS immunized mice (Figure 4B). To determine if our prime-and-trap vaccine can generate CS-specific liver-resident Trm, CS-tetramer-labelled CD8^+^ T cells were identified by either CD69^+^/KLRG1lo or CD69^+^/CXCR6^+^ expression. Both populations of CS-specific liver-resident Trm were larger in the 5-day prime-and-trap cohort and the RAS-immunized mice than in the 14-day prime-and-trap immunized mice (Figure 4C). This result is consistent with the observation that post-trap liver parasite burden is lower in the 14-day group, as shown above, resulting in less Trm generation.

**Figure 4.**
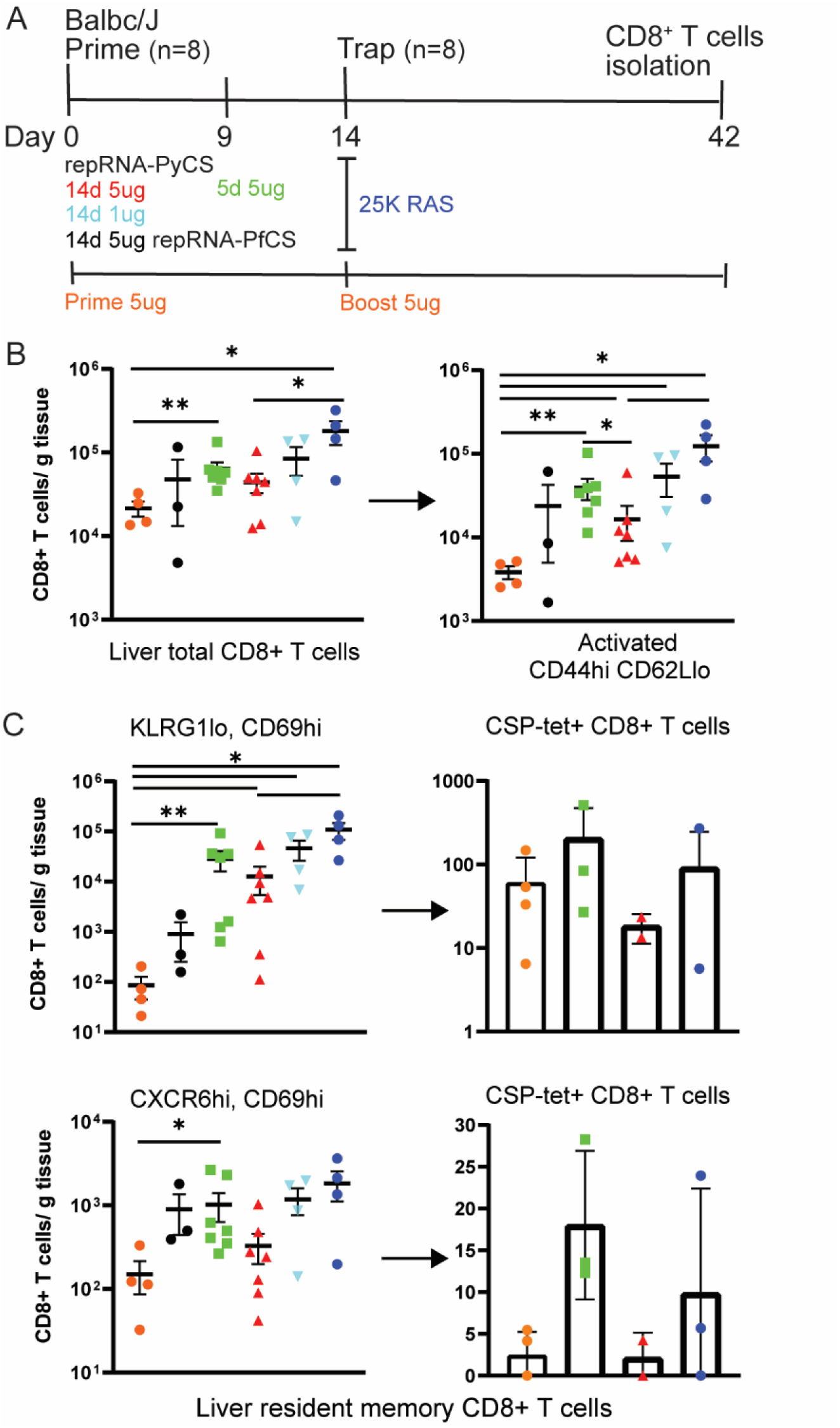
5-day prime-and-trap and trap-only (RAS) regimens of immunization induce higher-frequency liver Trm cells than 14-day prime-and-trap regimen. A) Schedule of immunization. 5ug 5-day (square, green data) or 5ug 14-day (triangle, red data) or 1ug 14-day regimen of repRNA-PyCS (inverted triangle, light blue data) prime followed by trap dose of 25,000 RAS (dark blue data), were tested in mice. Control cohorts are prime-boost repRNA-PyCS (orange data) or trap cohort (repRNA-PfCS +RAS, black data). B) Flow cytometric total CD8^+^ T cells and activated (CD44hi/CD62Llo) CD8^+^ T cells in perfused livers 28 days after the Trap dose of 25,000 RAS. C) Flow cytometric analysis of tetramer-stained, CS-specific CD8^+^ liver Trm cells (by CD69+ and either KLRG1lo (upper) or CXCR6+ (lower)). All error bars are SD of the mean. *p=0.05, **p=0.01, ***p=0.001 by Mann–Whitney two-tailed test. The n value represents total number of mice tested per cohort in two independent assays. Each data point represents an individual mouse and the bar represents the group mean with error bars representing the standard error of the mean. Asterisks represent significance as determined by the non-parametric two-tailed Mann–Whitney U test (*p=0.05, **p=0.01, ***p=0.001, ****p<0.0001).

### Efficacy of accelerated prime-and-trap repRNA-PyCS-RAS vaccination in BALB/cJ mice

To assess whether the immune responses induced by prime-and-trap immunization can confer sterile immunity, cohorts of mice immunized with LION/repRNA-PyCS as prime dose followed by RAS for the trapping dose (as described above in Figure 3) were challenged three weeks later with 1,000 freshly prepared WT Py spz delivered IV (Figure 5A, Supplemental Table 1). Upon challenge, 66% of trap-only and 80% of prime-and-trap mice were sterilely protected (Figure 5B; naïve mice were not protected). These results (66% vs 80%) are statistically equivalent. However, in the mice of the immunized cohorts that experienced breakthrough parasitemia, we saw a consistent 2- to 3-day delay in the onset of blood-stage parasitemia and a reduced peak load (Figure 5C). Two weeks post-challenge, sera were collected and total IgG levels were quantified by ELISA. The total IgG titer post-challenge was similar in all immunized cohorts (prime-and-trap vs trap alone), suggesting a recall of the CS-specific humoral immune response following spz challenge (Figure 5D).

**Figure 5.**
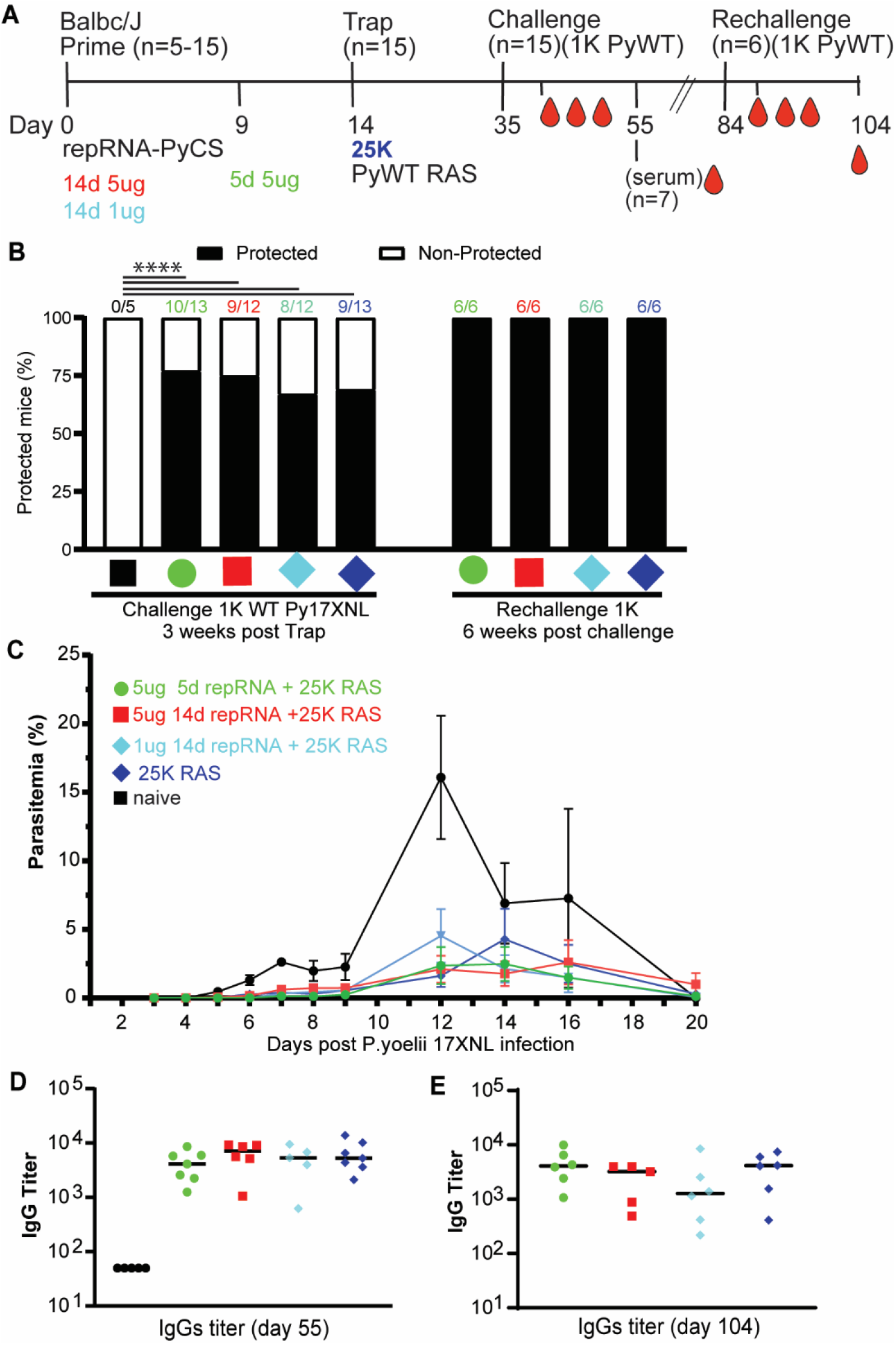
Efficacy of accelerated prime-and-trap immunization regimens in BALB/cJ mice. A) Immunization schedule. Prime dose of 5ug 5-day (green data) or 5ug 14-day (red data) or 1ug 14-day regimen of repRNA-PyCS (light blue data) followed by trap dose (dark blue data) of 25,000 RAS, were tested in mice. Three weeks later mice were challenged intravenously with 1,000 live spz isolated from infected mosquitos. Control cohorts are the trap cohort (RAS alone, dark blue data) and naïve cohort. Cohort showing partial protection were rechallenged with 1,000 live spz six weeks later. B) Protection post-challenge and re-challenge per cohort. Number of mice per cohort indicated above bar graph as protected/non-protected ratio. C) Parasitemia post-challenge of immunized mice cohorts. D) Sera from seven mice per cohort were harvested 20 days post-challenge (day 55) and CS-specific IgG titers were analyzed by ELISA. E) Final bleeds (six weeks post-rechallenge, day 104) of the four cohorts of mice showing full protection. The n value represents the total number of mice tested per cohort, in two or three independent assays. Each data point represents an individual mouse and the bar represents the group mean. All others statistical analyses were performed using a Mann-Whitney two-tailed test.

The four cohorts exhibiting partial sterile protection post-challenge were re-challenged 6 weeks after the first challenge (Supplemental Table 1). All were protected (Figure 5B) with high levels of circulating specific anti-CS IgG (Figure 5E). On the whole, the data from these replicate experiments are consistent with previous observations of limited protection after a single dose of PyRAS^2^, indicating that the prime-and-trap strategy is important for making the most of the LION/repRNA-primed responses.

### Sterile protection following same-day prime-and-trap repRNA-PyCS-RAS vaccination

As described above, accelerating the prime-and-trap vaccination from a 14-day to a 5-day immunization schedule reduced levels of circulating CS-specific antibodies (Figure 3B) and improved numbers of CS+ liver Trm (Figure 5) while retaining strong efficacy (Figure 4E). We next compared the protective efficacy in BALB/cJ mice of same-day and 5-day prime-and-trap regimens. Mice were vaccinated with 5 μg repRNA-PyCS and 25,000 RAS and challenged 3 weeks later by an IV dose of 1,000 freshly dissected infectious WT Py spz (Figure 6A, Supplemental Table 1). These cohorts were compared to a trap-alone (repRNA-PfCS + 25,000 RAS) cohort and an infectivity-control cohort. While both the trap cohort (3/5 mice) and control cohort (5/5 mice) rapidly developed high levels of parasitemia, parasitemia appeared in only 2/10 and 1/9 of the 5-day and 0-day prime-and-trap cohorts, respectively (Figures 6B-D). In the prime-and-trap cohorts, parasitemia appeared 7-8 days after infection and was cleared by day 16 (Figure 6B), whereas in both control cohorts the parasitemia appeared 4-5 days post-infection, persisted for 15 days, and was cleared at day 22 post-challenge as anticipated.

**Figure 6.**
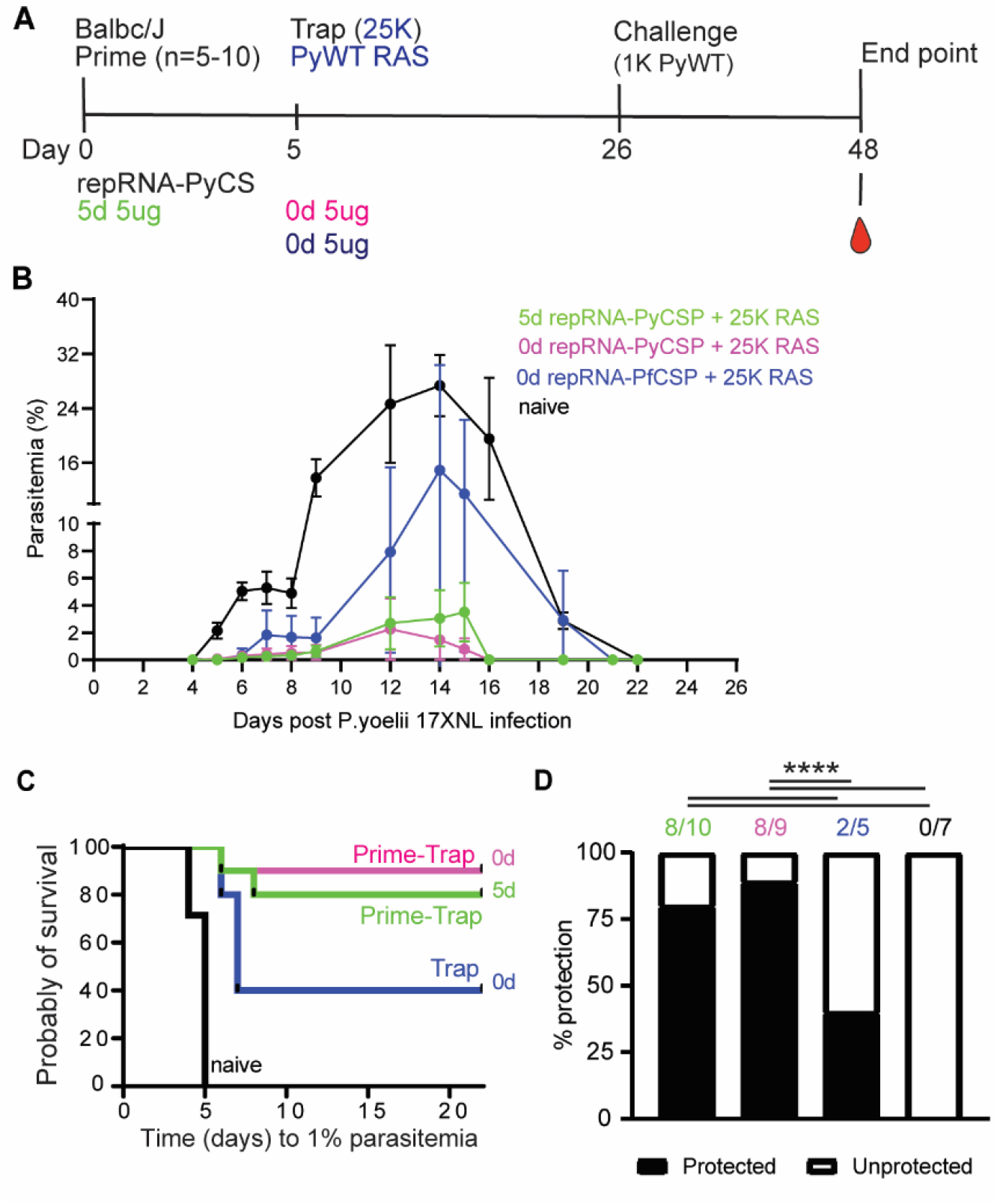
Immunogenicity and efficacy of a same-day prime-and-trap vaccine in BALB/cJ mice. A) Schedule of immunization. Prime-and-trap vaccine consisted of a 5ug 5-day regimen (green data) or 5ug same-day interval (pink data) of prime with repRNA-PyCS followed by trap dose of 25,000 RAS. Control cohort is 5ug same-day interval of a trap cohort (repRNA-PfCS +RAS, dark blue data). Three weeks later mice were challenged intravenously with 1,000 live spz isolated from infected mosquitos. B) Parasitemia post-challenge of all cohorts including the naive cohort (black data). C) Patency curves (>1% parasitemia) of mice post challenge. D) Protection post-challenge per cohort. Number of protected mice per cohort indicated above bar graph. ****p< 0.0001 by Fisher exact test. The n value represents total number of mice tested per cohort, in two independent experiments. Number of protected mice per cohort indicated above bar graph. ****p< 0.0001 by Fisher exact test. The n value represents total number of mice tested per cohort, in one experiment. All others statistical analyses were performed using a Mann Whitney two-tailed test *p=0.05, **p=0.01, ***p=0.001.

## DISCUSSION

An effective malaria vaccine will save many lives and lift the global burden of this disease. Among the next-generation approaches that build on the recent success of vaccines for COVID-19 prevention are mRNA vaccines delivered in nanoparticles ^36,37^. Our prime-and-trap approach combines two strategies that exhibit modest efficacy individually, but confer promising levels of protection when combined. This approach also has key logistical advantages over other current candidates. We summarize all the mouse immunization and challenge studies performed in this manuscript with Py wild-type parasites in Supplemental Table 1. We have shown that a homologous LION/repRNA-PyCS vaccine (prime-boost) is highly immunogenic at all doses evaluated, eliciting strong antibody responses when given in a 2-dose immunization regimen 2 weeks apart. Antibody levels are modest after the priming dose but increase subsequent to the booster dose. We have also demonstrated that homologous LION/repRNA-CS vaccination alone was insufficient to prevent blood-stage infection and did not provide protection in mice. To overcome these issues and enhance the practicability of vaccinating people in low-resource environments, our efforts have focused on minimizing the interval between injections, lowering dosages, and simplifying storage and preparation of the agents. We believe an effective vaccine based on a strategy such as that described herein, administered in a single clinic visit, will help to 1) enhance the logistics of the vaccination schedule, and 2) reduce the cost of goods by combining a repRNA priming dose and a single dose of attenuated spz.

Our results suggest that the priming dose of LION/repRNA-PyCS is highly immunogenic. We observed induction of a strong humoral response in BALB/cJ and C57Bl/6 mice when they were immunized on the 14-day, 5-day, or same-day regimens. We also observed a strain-specific CD8+ T-cell response in BALB/cJ mice. There was a 2-3-day delay in the onset of blood-stage parasitemia compared to the control groups. Compared with RAS-only immunization strategies, our prime-and-trap approach has the advantage of inducing both humoral and cellular immunities without compromising the generation of protective liver-resident CD8^+^ T cells.

Previous studies have shown that RAS vaccines can induce liver CD8^+^ Trm cells in animal models, but they can only achieve high levels of sterile protection in humans with three or more IV doses ^22,51,52^. The requirement for several IV doses compounds the time, cost, and logistical problems inherent to WO vaccines^29,30^. Others studies employing later-arresting GAP vaccines ^50,51,53^ have shown that presenting more antigens leads to improve protection over early-arresting GAP or RAS^18,54^. Likewise, immunization with wild-type (WT) WO spz administered under blood-stage drug chemoprophylaxis (known as CPS, (PfSPZ-CVac)^24,55^) allows full liver-stage development and can induce high levels of protection at reduced doses, although CPS involves co-administration of drugs with the vaccine, presenting practical and regulatory challenges^24^. These well-established immunizations with WO SPZ RAS ^56,57^, GAP ^18,50,53,58^, or PfSPZ-CVac^24,55^ are at least in part reliant on the generation of liver-resident memory CD8^+^ T cell (Trm) responses ^21^ ^33,59–61^.

In the present study, we show the contribution of CS-specific liver-resident memory CD8^+^ T cells induced by our prime-and-trap regime with LION/repRNA-PyCSP. Given that the initial dose of RAS administered to naïve individuals appears to be the most immunogenic and effective in inducing liver Trm cells, a vaccination strategy employing a single dose of RAS^34^ may be the most efficient and cost-effective means of using this valuable resource.

Production of Pf SPZ (RAS or WT) in mosquitoes is an expensive and tedious process. However notable progress in *in vitro* culture of Pf SPZ under current Good Manufacturing Practices has been made by Sanaria^®^. It is hoped that their cGMP mosquito-dissected spz, known as Pf SPZ vaccine (irradiated PfSPZ) or PfSPZ-CVac (PfSPZ used with chemoprophylaxis vaccination), will soon be available for human clinical trials ^3,22,62,63^. Current clinical trials involving injection of several doses of cryopreserved Sanaria PfSPZ have proven that large-scale implementation in endemic areas using liquid-nitrogen storage conditions is achievable ^29,30,64,65^. The combination of our repRNA/LION-based priming dose with a single dose of *in vitro*-produced spz may yield a highly effective, convenient, and cost-effective malaria vaccine.

Our findings reinforce the concept that CS remains one of the most immunodominant and protective antigens expressed by spz and suggest that its CD8^+^ T-cell epitope is involved in the protective effect against parasites in the liver of the BALB/cJ mice following RAS immunization ^66^. However, it is not clear that any single antigen will confer the robust and durable protection needed to eradicate malaria. Given the genetic diversity among parasite strains and the many variables in physiology, environment, and logistical capabilities involved in a broad vaccination campaign, it may be necessary to target multiple antigens from various stages of the parasite life cycle to eliminate malaria. A multi-stage malaria vaccine with high efficacy and durability is more likely to achieve eradication or major reductions in disease burden and transmission than a single-stage vaccine. With this in mind, we are currently evaluating the immunogenicity of repRNA presenting multiples antigens from various stages of the parasite life-cycle.

As seen during the ongoing SARS-CoV-2/COVID-19 pandemic, the use of RNA vaccines has numerous advantages over conventional vaccine approaches. These include the flexibility and the rapidity of the manufacturing processes, which proved important during the SARS-CoV-2/COVID-19 pandemic ^55^. Another advantage is the ease of production of the antigen-encoding RNA, which, when translated in host cells, yields the antigen that engages the host immune system through MHC class I and II pathways, resulting in T-cell responses and antibody production ^49^. Despite the advantages of RNA vaccines, however, the method of intracellular delivery of the RNA is also critical. The Pfizer/BioNTech and Moderna mRNA vaccines both employ LNPs to protect the mRNA from extracellular ribonucleases and facilitate cellular uptake via endocytosis ^39,40,67^. The mRNA-PfCSP LNP vaccine can provide from 40% to 80% protection in a lethal rodent infection model, depending on the dosage, schedule, and nucleoside-modified sequences of the mRNA, but the protection is short-lived^36^. Induction of antigen expression mediated by these mRNA-LNP vaccines is rapid but transient, yielding an immune response that is limited in magnitude and duration. While the mRNA-LNP vaccine approach is versatile and its efficacy is promising ^36^ more robust and durable antigen expression will be needed to reach efficacy objectives.

Our approach of combining repRNA and LION has several advantages over mRNA-LNP vaccines, including a reduced number of doses (2), the use of a single dose of the WO irradiated spz, a shorter (5-day or same-day) administration schedule, and simpler logistics ^36^. A further advantage of nanoparticle formulations as carriers for nucleic acid delivery is the possibility of optimization for specific target cells and tissue types. The balance of tissue targeting can be shifted by changing the nanoparticle composition and route of delivery (oral, subcutaneous, or IV) ^68^. One focus of our future work is direct targeting of the liver, in the hope of replacing spz administration with a more practical and cost-effective nanoparticle formulation. Studies to assess the efficacy and durability of our prime-and-trap approach over an interval of two months, to be extended in future work, are underway. In the current study, we used an intramuscular LION formulation that has been distributed in India under the product designation GEMCOVAC-19. This LION is composed of a stable cationic (DOTAP) squalene emulsion embedded in a hydrophobic oil phase, proven to be stable for weeks and highly immunogenic when injected intramuscularly ^44,45,69^. Of critical importance to any vaccine designed for use in a low-resource environment, GEMCOVAC-19 is manufactured as a lyophilized product and reconstituted prior to use, allowing long-term storage without the freezers and power requirements of other SARS-CoV-2 vaccines. Similarly, we expect to lyophilize our repRNA malaria vaccines to eliminate cold-chain requirements prior to reconstitution.

In summary, we have demonstrated a multi-component vaccination approach that concurrently induces humoral and T-cell immunities, using repRNA-CS formulated with LION nanoparticles and PyRAS targeting the liver. This prime-and-trap approach is currently being broadened in mouse studies to include antigens from other stages of malaria. We are also extending our protection studies to longer intervals to assess the durability of the observed efficacy. Our prime-and-trap approach is also being tested in non-human primate models in the US, potentially leading to immunogenicity and efficacy studies in a CHMI trial in the near future.

## ACKNOWLEDGMENTS

In memory of Dr Anil Ghosh.

We thank Tess Seltzer, Cecilia Kalhtoff, and Dr Alexis Kaushansky (Seattle Children’s Research Institute) for assistance and support of *Plasmodium yoelii*–infected mosquito production and dissection of the salivary glands. We thank Mint Laohajaratsang for her technical assistance and the vivarium staff of Bloodworks NW Center for assistance with the mouse facilities.

## AUTHORS CONTRIBUTIONS

M.A., J.H.E, A.P.K, B.W., S.C.M, S.G.R., J.W.D. conceived the research concepts. M.A., Z.M, J.H.E., A.P.K., B.W., M.J.S., S.C.M, S.G.R., J.W.D. designed this study. J.W.D secured the funding for this study. M.A., Z.M, K.H, M.L., A.S. M.J.S. and F.W., performed the experiments. M.A., Z.M, K.H., J.H.E., A.P.K., B.W., M.J.S., S.C.M, J.W.D participated in data analysis and interpretation. The manuscript was written by M.A. with revisions from all the authors.

## MATERIALS AND METHODS

### Vaccine design

The antigen chosen for our vaccine design is the full-length circumsporozoite (CS) protein from *Plasmodium*, as described in detail in Supplemental Figure 1. This antigen has a CD8^+^ T-cell epitope that is immunodominant and protective in the H2-K^d^ (BALB/cJ)-restricted genetic background.

### RNA Production and LION Formulation

Full-length CS coding sequences from *Pf*, *Pv*, and *Py* were cloned separately into a Venezuelan equine encephalitis (VEE) replicon vector (pT7-VEE-Rep). *In vitro* transcription was performed at 34°C using a T7 MEGAscript T7 Transcription kit (Invitrogen). RNA was purified via lithium-chloride precipitation, followed by capping with a capping kit (New England Biolabs) as described ^44^. RNA was further purified and stored at -80^0^ C until use. Denatured repRNAs were verified by electrophoresis in a 1% agarose gel. Briefly, 2 μg of each repRNA was denatured by glyoxal treatment (NothernMax-Gly, AM8551, ThermoFisher), and run in an agarose gel with NorthernMax-Gly gel prep/running buffer (AM8678, ThermoFisher). Ethidium bromide was premixed into the running buffer and the gel image was analyzed by a Biorad gel docXR+ Imaging system.

To protect the RNA replicons from degradation, we combined them with LION nanoparticles obtained from HDT Bio ^44^. LION nanoparticles consist of a hydrophobic squalene oil core stabilized with Tween 80, Span 60, and the cationic lipid DOTAP. The oil (squalene, span 60 and DOTAP) and aqueous (Tween 80 in 10 mM sodium citrate) phases were homogenized using an L5M-A high-shear mixer (Silverson) and further processed by passaging through a microfluidizer to achieve an average hydrodynamic diameter of 60 nm and polydispersity index of 0.2 by dynamic light scattering. The microfluidized LION was terminally filtered with a 200-nm pore-size polyethersulfone filter and stored at 2° to 8°C.

### Cells lines

To qualify the vaccine candidates *in vitro*, BHK cells (American Type Culture Collection) were transfected with repRNA or mock-transfected using OptiMEM (Gibco) and Expifectamine transfection kit (ThermoFisher). Cells were scraped off and lysed with RIPA buffer 24-48 hours later, and lysates were analyzed by SDS–polyacrylamide gel electrophoresis and Western blot.

### Western blots

Cells lysates were analyzed by Western blot after transfer to nitrocellulose membranes. For detection, anti–rabbit polyclonal anti-CSP (Py, Pf, or Pv) antibodies (Pocono) were used (1/1,000) followed by goat anti-rabbit IgG (H+L). An alkaline phosphatase-linked secondary antibody was used (Invitrogen, T2191) (1/10,000).

### Ethics Statement

Animal studies were performed according to the regulations of the Institutional Animal Care and Use Committee of Bloodworks Northwest, and approval was obtained from this committee. Our studies meet the standards of the Guide for the Care and Use of Laboratory Animals and applicable Bloodworks Northwest policies and procedures. Bloodworks Northwest has an approved Animal Welfare Assurance (#A4659-01, D16-00862) on file with the NIH Office of Laboratory Animal Welfare (OLAW).

### Mice

Female BALB/cJ mice, six to eight weeks old, were purchased from The Jackson Laboratories (Bar Harbor, ME, USA). Mice were maintained under pathogen-free conditions in animal facilities and were fed autoclaved food ad libitum. Mice were housed and cared for in standard IACUC-approved animal facilities at Bloodworks Northwest and used in compliance with IACUC-approved protocols.

### LION/repRNA vaccination

For all LION/repRNA vaccines, 5 μg of RNA was mixed with LION at a nitrogen:phosphate (N:P) ratio of 15 and injected IM into mice using a total of 50 ul (25 ul in each leg). A two-vial formulation method was performed as described ^44^. Our immunization protocol and timeline are given in each respective figure.

### Sporozoite isolation, vaccination, and challenge

Wild-type Py (17XNL strain) spz were prepared by cyclical transmission in BALB/cJ mice and *Anopheles stephensi* mosquitoes at the Seattle Children’s Center for Global Infectious Disease Research Insectary (Seattle, WA, USA). Female 6- to 8-week-old Swiss Webster (SW) mice were injected with blood-stage Py 17XNL WT parasites to begin the growth cycle and used to feed female *Anopheles stephensi* mosquitoes. At day 15 after blood meal, salivary-gland spz were isolated and harvested as previously described ^70^. RAS were generated by exposure to 10,000 rads using an X-ray irradiator (Rad-Source, Suwanee, GA, USA). RAS were resuspended in 100 μL Schneider and administered to the mice through tail-vein injection. Infectious spz for challenge were prepared in an equivalent manner but without irradiation. All experimental and control mice were challenged with live *Py* 17XNL spz. A summary of vaccination and challenges experiment is included in Supplemental Table 1.

### Liver lymphocyte isolation and flow cytometry

Livers were perfused with 10 ml PBS/2 mM EDTA by injection into the portal vein, with outlet drainage via the inferior vena cava, and mashed into a single-cell suspension. Intrahepatic lymphocytes were isolated as previously described^34,35^. Final pellets were resuspended in 150 μl 1x MACs buffer and transferred to a 96-well plate for blocking and staining prior to flow cytometry. All antibody characterizations and flow-cytometry analyses were performed as previously described^28,35^, using a live/dead dye (Zombie NIR Fixable Viability Kit, BioLegend) to enable exclusion of dead cells from downstream analysis. In brief, liver lymphocytes were treated with an Fc block (anti-CD16/32, clone 2.4G2; BD Biosciences) and live/dead dye for 30 min, stained for 45 min (with antibody cocktail as described^35^), and fixed for 20 min (Cytofix/Cytoperm reagent; BD Biosciences). Cells were gated for CD8+ T cells (CD3e+, B220-, CD4-), CD44hi by CD62Llo, then assessed by either KLRG1lo by CD69hi or by CXCR6hi by CD69hi. Antigen specificity was then assessed by PyCSP-tetramer (SYVPSAEQI-specific H2-Kd tetramer, National Institutes of Health Tetramer Core) conjugated to streptavidin-allophycocyanin (ProZyme) per standard protocols. Cell count per gram of tissue was calculated based on a known concentration of counting beads per sample to normalize data. Flow cytometry was performed on an LSR II (BD Biosciences), and data analyzed with FlowJo version 10.7.1 (BD Biosciences).

### *Ex vivo* IFNγ ELISPOT

Spleens were harvested and splenocytes separated from BALB/cJ mice 28 days post-immunization. A total of 1×10E5 splenocytes were combined with SYVPSAEQI peptide (1 mg/ml final) (Genemed Synthesis) for murine IFNγ ELISPOT (eBioscience), cultured for 18 h at 37°C, and developed following manufacturer guidelines. The percentage of antigen-specific T cells was calculated based on the spot-forming units counted in each well divided by the total number of splenocytes applied to each well.

### Blood stage and liver burden

Breakthrough to blood-stage patency was assessed by Giemsa-stained thin blood smear starting at day 4 after challenge and ending at day 21, at which time a negative smear was attributed to complete protection. Liver burden was detected by qRT-PCR from harvested liver 44 hr post-challenge ^34,71^.

Mice immunized with Pf- or Pv-repRNA-CS and challenged with live Py spz were used as controls. Sterile protection was defined as being blood-smear negative. The Kaplan-Meier curves illustrate the time to developing a parasitemia during days 4–21 after challenge with 17XNL strain Py live spz.

### qRT-PCR

Total RNA was extracted from Py-infected livers using TRIzol reagent (Thermo Fisher Scientific) and treated with Turbo DNase (Ambion). cDNA synthesis was performed using a SuperScript III Platinum two-step qRT-PCR kit (Thermo Fisher Scientific). Specific PCR primers (as listed below) were used to amplify *Py* 18S rRNA and GAPDH (housekeeping) gene from cDNA derived from mouse liver and spleen. The primers used for amplification of 18S rRNA from cDNA were 18S-fwd: (GGGGATTGGTTTTGACGTTTTTGCG) and 18S-rev: (AAGCATTAAATAAAGCGAATACATCCTTAT). Mouse GAPDH was amplified with cDNA using gapdh-fwd: (CCTCAACTACATGGTTTACAT) and gapdh-rev: (GCTCCTGGAAGATGGTGATG) primers. All qRT-PCR amplification cycles were performed at 95°C for 30s (DNA denaturation) and 60°C for 4 min (primer annealing and extension). Samples are run in triplicate. Results are expressed as the difference ΔCT in threshold cycle number between the average of CT value of the Py 18S parasite gene and the average of CT value of the GAPDH house-keeping gene, so a high ΔCT represents a low parasite burden.

### ELISA

MaxiSorp plates were coated with 100 μL CS (Py, Pf, or Pv) peptide at 1μg/ml in PBS. Plates were then washed with PBS + 0.05% Tween 20 (PBS-T) and blocked with 1% BSA in PBS-T for 2 hrs at RT. Murine serum samples were plated at a dilution of 1:50 in PBS-T +0.1% BSA, serially titrated 1:3 for 6 wells, and incubated for 2 hrs at RT or overnight at 4°C. Following washing steps, plates were incubated with secondary antibodies diluted 1:5000 in PBS-T + 0.1% BSA for 1 hr at RT. After a second wash, 100 μL TMB was added per well and incubated 5-10 minutes before stopping with 50 μL 1N sulfuric acid.

### Statistics

Comparisons of ELISA groups or flow-cytometry cell counts were done using the non-parametric two-tailed Mann–Whitney U test (*p=0.05, **p=0.01, ***p=0.001, ****p<0.0001). ELISPOT assay comparisons were done by unpaired, two-tailed Student’s t tests. Statistical significance between groups of mice for their spleen or liver burden qRT-PCR was evaluated using one-way ANOVA followed by the Kruskal-Wallis test and Dunn’s multiple-comparisons test (* p<0.05, ** p<0.005). Protection data were evaluated using Fisher’s exact test. All groups were compared against the prime-boost cohort or the trap-alone cohort (repRNA-PfCS or repRNA-PvCS for priming, and RAS for trapping dose; **** p< 0.0001). Error bars are SEM of the mean with individual mouse samples shown. Statistical significance was defined as p<0.05 using Prism Graph-Pad 9.4.1 Software (San Diego, CA).

