## Supplemental for "Accelerated prime-and-trap vaccine regimen in mice using repRNA-based CSP malaria vaccine"

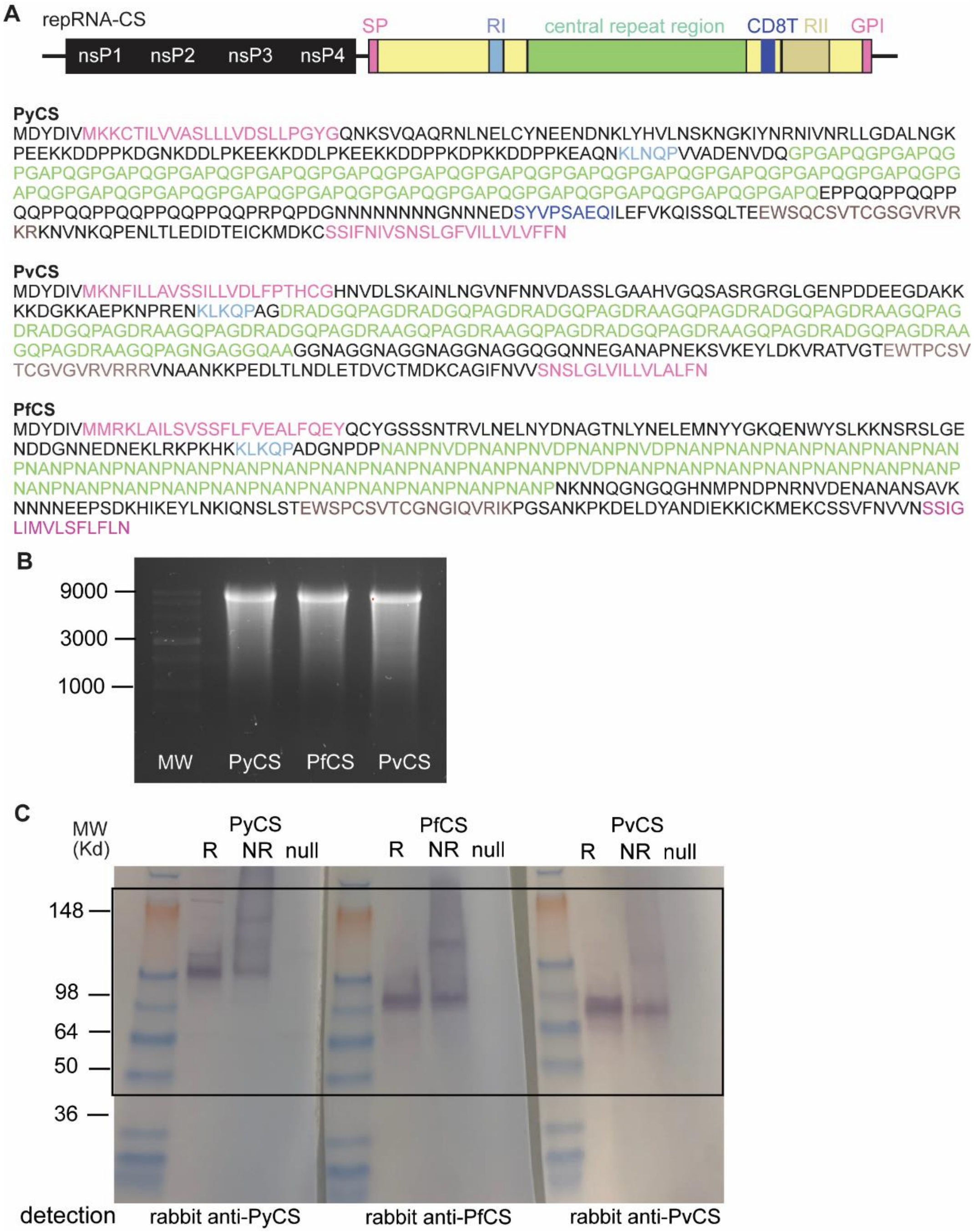


**Supplemental Figure 1. CS replicon and protein sequences.** A) Schematic of the replicon encoding the full length of either *P. yoelii*, *P. vivax,* or *P. falciparum* circumsporozoite (CS) protein including the signal peptide (SP) at the N-terminus, the region I (RI), the entire central repeat region, the CD8$^{+}$ T cell epitope, the region II (RII) up to the GPI anchor signal in C-terminal. The replicon was cloned into an alphavirus replicon encoding the four nonstructural protein (nsP1 to nsP4) genes of the Venezuelan equine encephalitis virus (VEE) strain TC-83. For each CS cloned into the replicon, the full protein sequence is indicated. B) Northern denaturing agarose gel of repRNA-PyCS, -PfCS, and -PvCS. C) Uncropped western blot from Figure 1A showing protein expression.

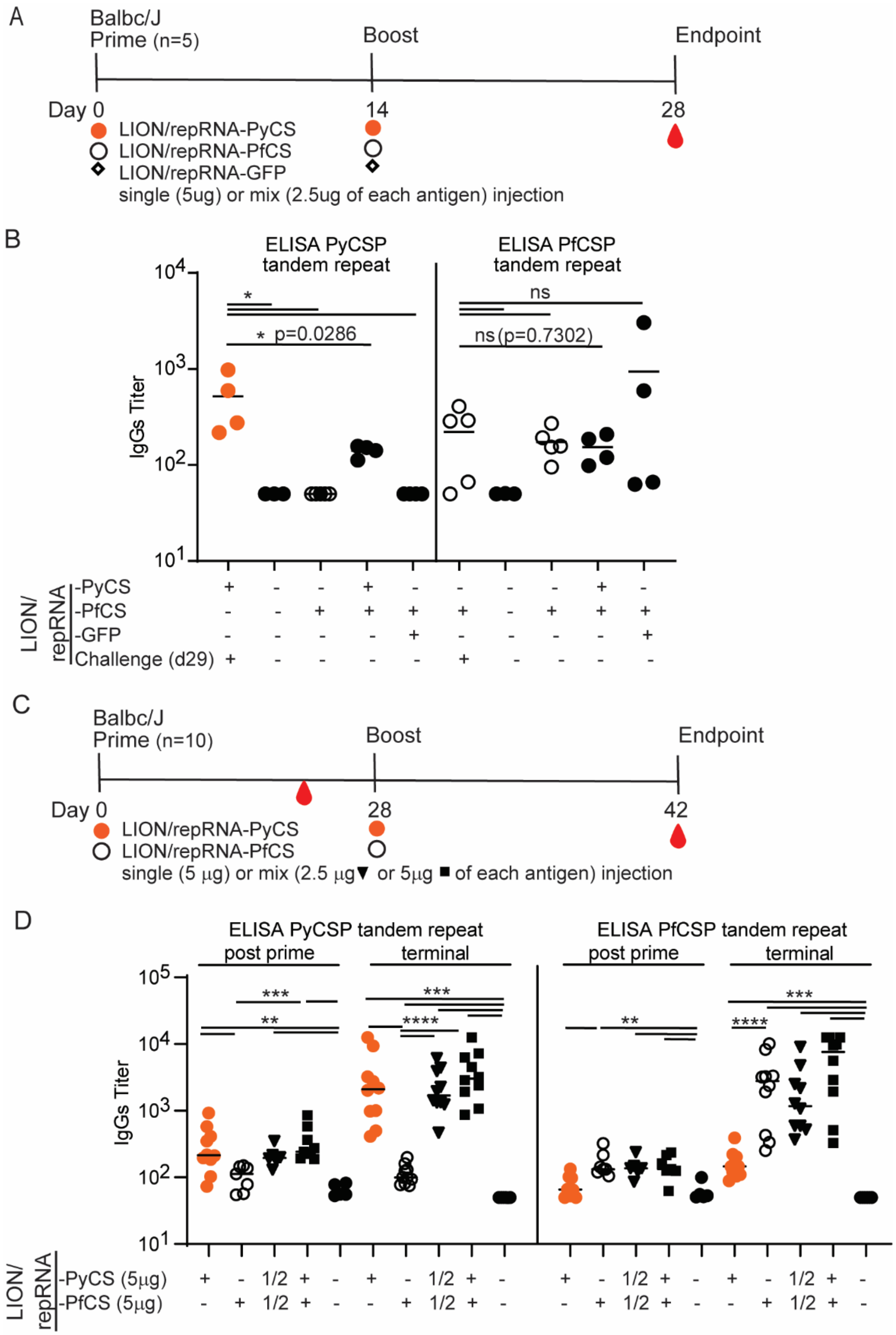


**Supplemental Figure 2. Immunogenicity in BALB/cJ of repRNA-CS formulated with LION.** A) Immunization schedule for the 14-day Prime-Boost with either LION/repRNA-PyCS, -PfCS. or -GFP used as control replicon. n=5 mice per cohort in one experiment. B) Mice were either immunized with a single antigen (5μg each) or two antigens mixed (2.5μg each). Final bleeds were collected two weeks post-boost at endpoint and immune responses were analyzed by ELISA against their corresponding CS tandem repeat regions. C) Immunization schedule for the 28-day Prime-Boost with either LION/repRNA-PyCS, or -PfCS. . n=10 mice per cohort in one experiment. D) Mice were either immunized with a single antigen (5μg each) or two antigens mixed (2.5μg each for a total of 5μg, labelled as 1/2) or two antigens mixed (5μg each for a total of 10μg). Final bleeds were collected four weeks post-boost at endpoint and immune responses were analyzed by ELISA against their corresponding CS tandem repeat regions.  Each data point represents an individual mouse, and the bar represents the group mean. Asterisks represent significance as determined by non-parametric two-tailed Mann–Whitney U test (*p=0.05, **p=0.01, ***p=0.001, ****p<0.0001).

**Supplemental Table 1:**

Summary of mice immunization and challenge studies with *P.yoelii* wild-type parasites.

| Immunization | | | | | Challenge | | | | | |
|---|---|---|---|---|---|---|---|---|---|---|
| BALB/cJ mice | PRIME dose (LION/ repRNA) | Route | 2nd dose (day given) | Route | IV dose (week given) | Blood-stage patency | Protected/ Challenged | % protection | p-value | Linked to |
| 15 | 5ug PyCSP | IM | 5ug PyCSP (day 14) | IM | 1000 WT SPZ (week 3) | 4 days | 0/13 | 0 | | Figure 2 |
| 7 | 5ug PfCSP | IM | 5ug PfCSP (day 14) | IM | 1000 WT SPZ (week 3) | 4 days | 0/5 | 0 | | |
| 7 | 5ug PvCSP | IM | 5ug PvCSP (day 14) | IM | 1000 WT SPZ (week 3) | 4 days | 0/7 | 0 | | |
| 12 | 5ug PyCSP | IM | 25000 RAS (day 14) | IV | 1000 WT SPZ (week 3) | 7 days | 9/12 | 75%* | <0.0001 | Figure 5 |
| 13 | 5ug PyCSP | IM | 25000 RAS (day 5) | IV | 1000 WT SPZ (week 3) | 7 days | 10/13 | 77%* | <0.0001 | |
| 12 | 1ug PyCSP | IM | 25000 RAS (day 14) | IV | 1000 WT SPZ (week 3) | 8 days | 8/12 | 67%* | <0.0001 | |
| 13 | - | - | 25000 RAS | IV | 1000 WT SPZ (week 3) | 6 days | 9/13 | 69%* | <0.0001 | |
| 5 | - | - | - | - | 1000 WT SPZ | 5 days | 0/5 | 0% | | |
| | | | | | *Rechallenge | | | | | |
| | | | | | 1000 WT SPZ (week 6) | none | 6/6 | 100% | | |
| | | | | | 1000 WT SPZ (week 6) | none | 6/6 | 100% | | |
| | | | | | 1000 WT SPZ (week 6) | none | 6/6 | 100% | | |
| | | | | | 1000 WT SPZ (week 6) | none | 6/6 | 100% | | |
| 10 | 5ug PyCSP | IM | 25000 RAS (day 5) | IV | 1000 WT SPZ (week 3) | 6-7 days | 8/10 | 80% | <0.0001 | Figure 6 |
| 9 | 5ug PyCSP | IM | 25000 RAS (day 0) | IV | 1000 WT SPZ (week 3) | 5 days | 8/9 | 89% | <0.0001 | |
| 5 | 5ug PfCSP | IM | 25000 RAS (day 0) | IV | 1000 WT SPZ (week 3) | 6-7 days | 2/5 | 40% | <0.0001 | |
| 7 | - | - | - | - | 1000 WT SPZ | 4 days | 0/7 | 0 | | |

* partially protected mice were rechallenged 6 weeks post-challenge
